# Interpreting mammalian evolutionary constraint at synonymous sites in light of the unwanted transcript hypothesis

**DOI:** 10.1101/2024.04.23.590689

**Authors:** Matthew J. Christmas, Michael Dong, Jennifer R. S. Meadows, Sergey V. Kozyrev, Kerstin Lindblad-Toh

## Abstract

The unwanted transcript hypothesis presents a potential explanation for cryptic evolutionary constraint at synonymous sites in species with low effective population sizes, such as humans and other mammals. Selection for higher GC content and against mutations that alter splicing in native transcripts is predicted to shape synonymous site content and protect against unwanted transcripts. Here, we interpret mammalian synonymous site constraint in this context. Utilising the largest alignment of 240 placental mammal genomes and single-base resolution constraint scores, we show that 20.8% of four-fold degenerate sites are under significant constraint across mammals. There is a strong bias for guanine (G) and cytosine (C) at constrained sites, marked constraint near splice sites, and variation in human populations shows a bias against mutations that reduce synonymous site GC content. We find evidence for higher constraint on four-fold degenerate sites in species with small historic effective population sizes and high young transposable element genome content. Genes enriched for synonymous site constraint, including those forming CpG sites, are tightly regulated and integral to organismal viability through their involvement in embryo development and transcriptional regulation.

## Introduction

Evolutionary constraint, or the restriction of sequence evolution over time due to strong purifying selection, is evident in functional regions of genomes. In protein coding genes, constraint is largely due to purifying selection acting on mutations that alter the amino acid sequence and disrupt protein function. As a result, mutations at synonymous sites that have no effect on the amino acid specified were originally considered as evolving neutrally^1^. It is now well established that selection can act on synonymous sites, particularly in fast-growing species^2^ and species with large effective population size (*Ne*), where efficient selection can optimise codon use to maximise translational velocity and accuracy^3,4^. However, substantial constraint on synonymous sites is also observed in the genomes of slower growing and low-*Ne* species, such as humans and other mammals, where evidence for codon usage bias due to translational selection is lacking or absent^5^.

Estimates of the amount of constraint at synonymous sites in mammals vary. A comparison of 1,572 mouse-rat orthologous genes detected purifying selection at synonymous sites in 28% of the genes^6^. Genome-wide estimates of constraint identified that ∼25% and ∼11% of four-fold degenerate (4d) sites are under constraint in hominids and murids respectively, based on human-chimpanzee and mouse-rat comparisons^7^. Beyond mammals, significant constraint at synonymous sites likely independent of translational selection has been identified in birds, with lineage-specific estimates of constraint on 4d sites between 24% and 43%^8^. In invertebrates, 22% of 4d sites in *Drosophila melanogaster* are estimated to have evolved under strong selective constraint, seemingly independent from translational optimisation, splicing enhancers, or nucleosome positioning^9^. In *D. melanogaster*, genes enriched in strongly constrained 4d sites are functionally important, with many involved in organism development^9^. Constraint at synonymous sites therefore seems to be a ubiquitous phenomenon across animals.

There is evidence for selection on codons due to translational optimisation^10^, protein folding^11,12^, accurate splicing^13,14^, and mRNA stability^15^ in mammals^16^. Additionally, overlapping regulatory features such as transcription factor binding sites^17^, being located within the binding sites of microRNAs or RNA-binding proteins^18^, or within upstream open reading frames (uORFs)^19^, may bring secondary functions to 4d sites. Selection for CpG dinucleotides, the site of DNA methylation in mammals, in certain transcription factors and developmental genes is also thought to lead to constraint at synonymous sites^20^. However, the selection pressures acting on the majority of synonymous sites in mammals largely remain elusive.

In vertebrate genomes, a pattern of high guanine (G) and cytosine (C) content at third codon positions is in contrast to a genomic background of low GC content due to mutational bias and is suggestive of selection^21^. Recently, the ‘unwanted transcript hypothesis’ (UTH) has been presented as a novel explanation for synonymous site GC bias and constraint in humans and other species with low *Ne*^22^. It proposes that selection on synonymous sites aids the retention of native, functional transcripts and the identification and removal of unwanted, spurious transcripts. Around 62% of the human genome is transcribed, yet only 1.22% of the genome encodes protein-coding exons^23^. Much of the transcription is likely to be spurious, generated from the integration of novel sequences into the genome such as from viruses and transposable elements (TEs), leading to costly transcripts whose removal is advantageous^22^. In species where *Ne* is low, unwanted transcripts may accumulate due to inefficient selection against their mildly deleterious effects^24^.

The UTH posits that much of the selection on synonymous sites is to reduce the production or improve the quality control of unwanted transcripts^22^. Base content of synonymous sites can increase the signal that a transcript is native and differentiate it from randomly transcribed or newly arrived sequences, such as from a retrovirus. In mammals, which generally have GC-poor genomes but show relatively GC-rich coding sequences^21^, high GC content can indicate that a transcript is likely functional and set it apart from spurious transcripts generated from AT-rich intergenic regions, providing a potential explanation for GC-rich synonymous sites. Human native transcripts are generally GC-rich, CpG poor (due to hypermutability of methylated CpGs), contain introns, and have short exons (80% are <200 bp^25^). Transcripts that are AT-rich, have high CpG content, are intronless, and contain long exons are therefore most likely to be spurious and will be targeted for removal. In agreement with this, it has been shown that higher GC content at synonymous sites of transgenes in mammals leads to increased protein levels related to increased nuclear export^26–28^, and intronless retrogenes tend to evolve higher GC content at 4d sites (GC4) than their intron-containing parent genes^26^. Selection on synonymous sites can therefore act to both reduce the production and increase the removal of spurious transcripts by ensuring accurate splicing and aiding in their discrimination respectively.

In this study, we utilised the largest alignment of 240 placental mammal species genomes, produced as part of the Zoonomia project^29^, to comprehensively characterise synonymous site constraint in mammals. We identified shared synonymous sites from the alignment, extracted their base content, and used single-base resolution phyloP scores generated in Zoonomia^30^ to characterise the landscape of mammalian synonymous site constraint. We provide an assessment of whether the constraint observed at synonymous sites fits with the UTH predictions of high GC content, low CpG content, and high constraint at splice sites. Additionally, we test whether variation in synonymous site GC content across the mammal species relates to *Ne* and a proxy for unwanted transcript burden. We also tested how much of the observed synonymous site constraint can potentially be explained by being located within regulatory features, such as transcription factor (TF) binding sites. This work provides the most comprehensive characterisation and assessment of mammalian synonymous constraint to date and advances our understanding of the selection pressures acting on synonymous sites in placental mammals.

## Results

### Constraint at 4d sites and GC-bias

We identified all codons containing 4d sites in the human genome using Gencode annotation v.39^31^ and taking the canonical transcript per gene (n = 19,386), revealing a total of 5,278,470 4d sites. Many of these are unlikely to be 4d sites in other mammal genomes due to, for example, differences in the gene content among genomes. We therefore used the Zoonomia alignment to identify codons where >85% of species were aligned and >95% of the aligned codons contained a 4d site (see methods). This resulted in a set of 2,621,118 4d sites distributed over 17,394 transcripts that formed the basis of our analysis (supplementary data file 1). We then extracted the base content of each 4d site within each aligned genome (at least 204 species) and used the single-base phyloP scores generated in Zoonomia as estimates of constraint^30^. These scores range from -20 to 8.9, with negative scores indicating accelerated evolution, scores close to 0 indicating neutral evolution and positive scores indicating constrained evolution. A significance threshold for constraint was previously established for this dataset using a 5% false discovery rate (FDR), where sites with a phyloP score ≥ 2.27 are under significant constraint^30^. Overall, we see significant constraint at 20.8% of 4d sites (table 1). In comparison, 29.4% of three-fold (3d), 36.6% of two-fold degenerate (2d) and 74.1% of nondegenerate sites are under significant constraint at this threshold. Synonymous sites with a phyloP score at the 5% FDR threshold have a mean of 230 species aligned (s.d. = 9.16) and a mean of 188.5 species sharing the same base (s.d. = 37.8). We observe 4,906 4d sites (0.19%) that are fixed for the same base across all the genomes.

**Table 1.**
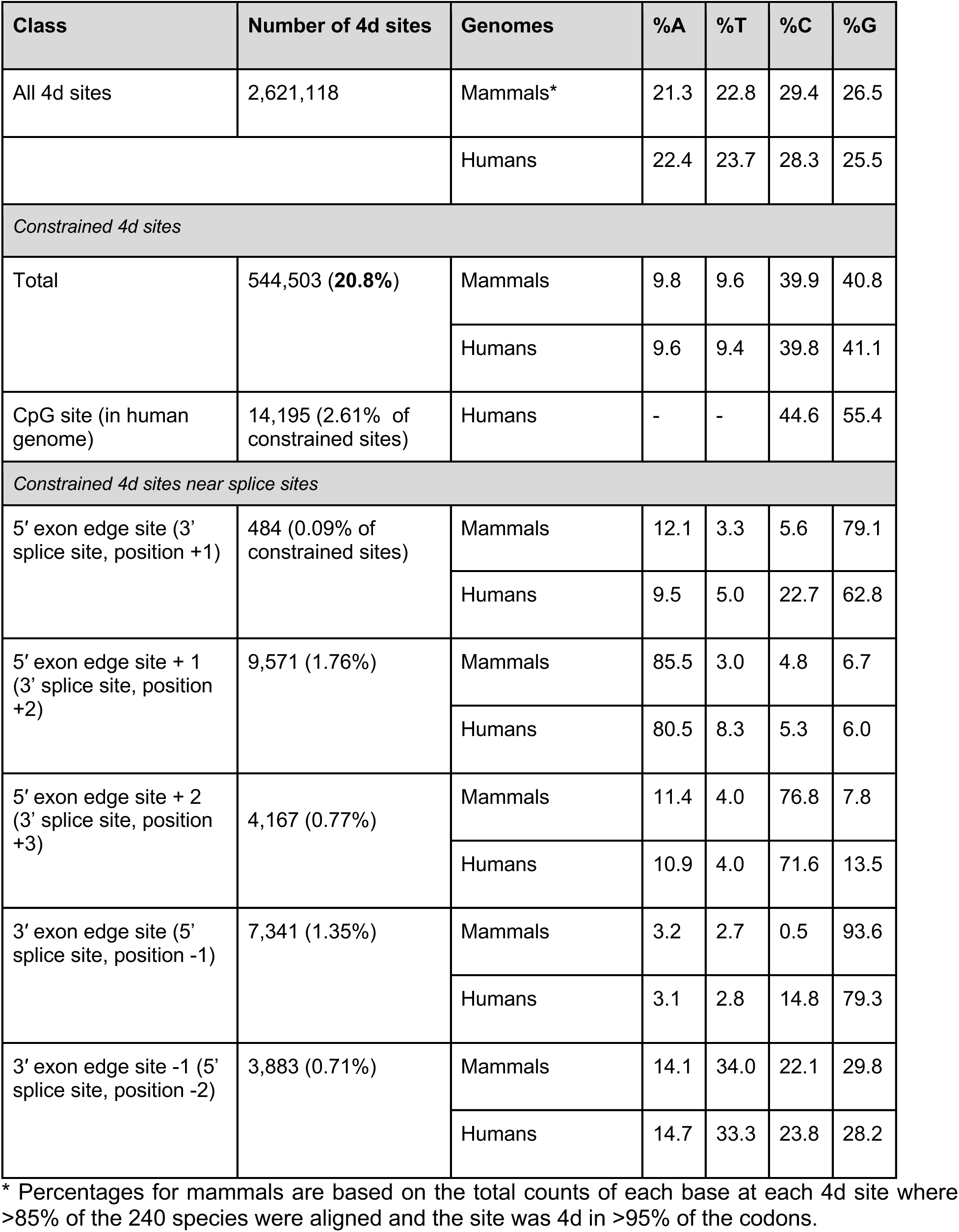
Summary of constraint at four-fold degenerate sites in mammals.

There is a strong bias for greater GC content at constrained 4d sites in mammals (Table 1), with only the hispid cotton rat genome (*Sigmodon hispidus*, Order: Rodentia) having a GC4 content below 50% (49.5%; Fig.1A). The common shrew (*Sorex araneus*, Order: Eulipotyphla) has the highest 4d GC content at 64.3%. For sites below the significant constraint threshold (phyloP < 2.27), observations of each base across the genomes are very similar (A = 24.2%, T = 26.0%, C = 26.7%, G = 23.1%). However, 80.7% of bases at constrained 4d sites are G or C (Table 1). A strong skew towards higher GC content at 4d sites under higher constraint is seen across the mammalian genomes generally (Fig. 1B-D). For any one 4d site under constraint, the base content is generally skewed towards a particular base in all mammals (Fig. 1E-F), which suggests selection for a certain base at a site, rather than a general increase in GC content.

**Figure 1.**
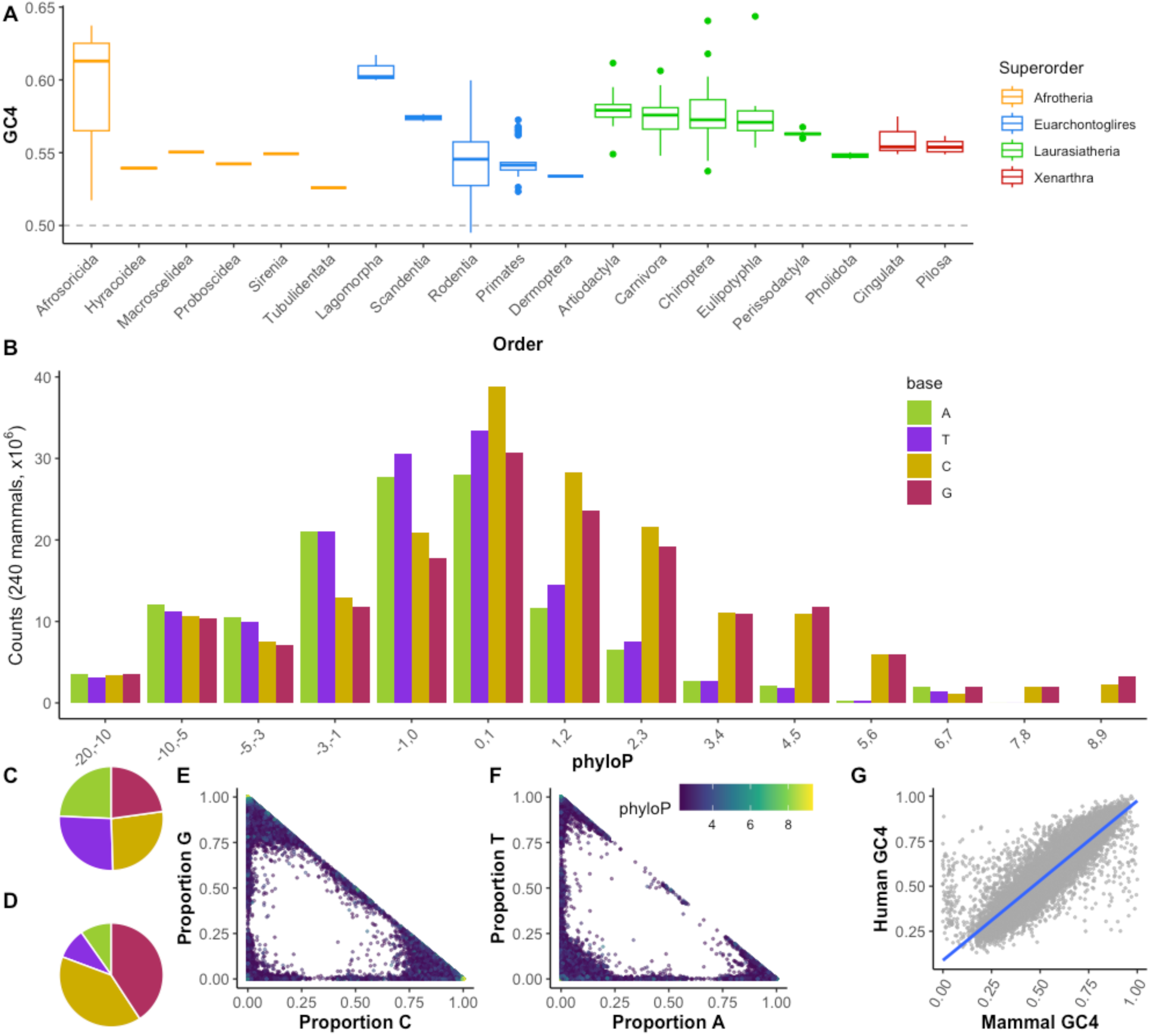
Constraint at four-fold degenerate sites in mammals is heavily GC-biased. (A) GC content of 4d sites across the species in Zoonomia. Only one species (the hispid cotton rat, *Sigmodon hispidus*, Order: Rodentia) has less than 50% GC at 4d sites (49.5%). (B) The number of each base at 4d sites across 240 placental mammal genomes, binned by phyloP score, where more positive scores indicate stronger constraint. Pie charts showing the proportion of each base seen at 4d sites across the mammal genomes for sites (C) phyloP < 2.27 and (D) phyloP ≥ 2.27. At constrained 4d sites (phyloP ≥ 2.27), we observe a general bias towards C or G at GC sites (E) and a bias towards A or T at AT sites (F) among the mammal genomes, suggesting selection for a particular base rather than an indiscriminate preference for C and G at 4d sites. (G) GC4 content of human transcripts and summarized across the mammal genomes shows a strong positive correlation (Pearson’s R = 0.90, p < 2.2×10^-16^). Trend line was calculated using a linear model.

CpGs are generally depleted in mammalian genomes due to the hypermutability of methylated CpGs, except for the CpG islands in the promoter regions of some genes^32^. Among all 4d sites, 10% of them form CpGs in the human genome. Constraint at 4d sites in human CpGs is significantly less than at other 4d sites, reflecting the hypermutability of these sites (n = 265,111 and 2,357,007, mean phyloP = -3.98 and 0.28 at 4d sites in and not in CpGs respectively; two sample t-test, t = 441.23, d.f. = 293,603, p < 2.2×10^-16^). However, 14,195 are under significant constraint (2.6% of constrained sites), which likely relates to selection for epigenetic regulation of certain genes via methylation^20^.

### Codon specific GC biases

Transcript GC4 content correlates strongly between humans and across the mammal genomes generally (Fig. 1G; Pearson’s R = 0.90, p < 2.2×10^-16^). The major allele in humans (identified from the genotypes of 138,922 individuals in the TOPMed dataset^33^) matches the most common base across the mammal genomes at 79.5% of all 4d sites, rising to 90.6% at constrained 4d sites. We generated base counts from the human genome across each of the codons containing 4d, 3d and 2d sites to assess whether GC bias is seen for all codons. Constrained 4d sites show a strong GC bias across all codons, with a minimum GC4 content of 59.5% (proline codons) and a maximum of 90.5% (valine codons; Fig. 2A). Differences in base proportions at 4d sites in the codons of each amino acid reflect further restrictions on base content. For example, in codons where the second base is cytosine (codons for alanine, proline, serine and threonine), 4d sites are rarely guanine, which would generate CpG sites. Indeed, 67% of constrained 4d sites in humans are G, and only 5% are C, when the subsequent base is G. In glycine codons (GGN), high cytosine content is observed at 4d sites (17% A, 61% C, 16% G, 7% T), likely due to selection against GGG codons that can lead to the formation of G-quadruplex structures in mRNA^34^. Again, if the subsequent base is G then 4d C content is low as this would generate CpG sites (38% A, 16% C, 32% G, 13% T). We also see strong biases for GC at constrained 3d and 2d sites in all codons except those for arginine. Constrained synonymous sites in all other codons are at least 72% C or G, with 74% C at constrained 3d sites in the codons for isoleucine (Fig. 2B,C). For unconstrained sites, we do not observe a general bias for higher GC content at synonymous sites (Fig. 2D-F).

**Figure 2.**
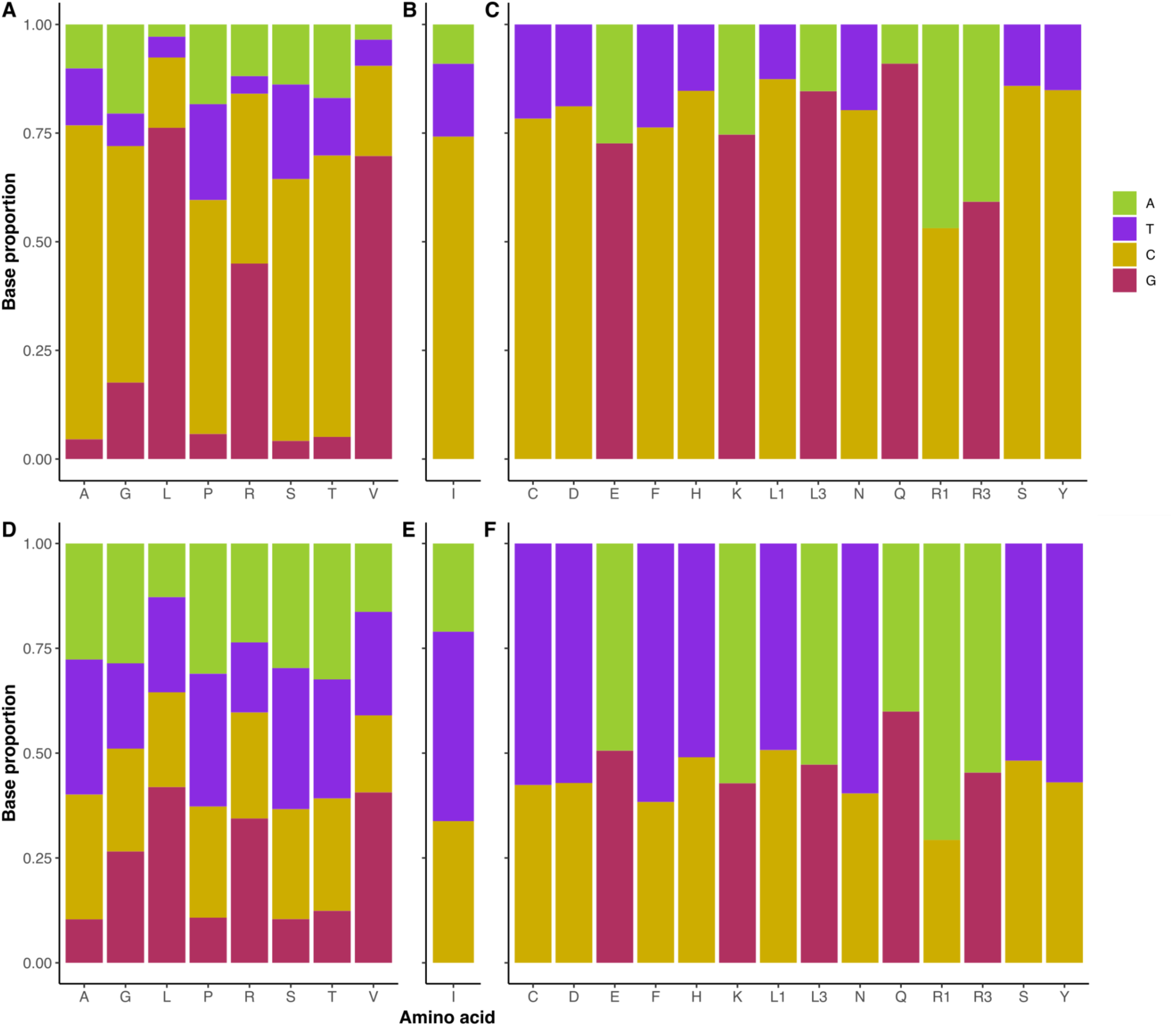
Differences in G and C content at constrained four-fold degenerate sites likely reflect restrictions on codon usage. Base content of significantly constrained (A) 4d, (B) 3d, and (C) 2d synonymous sites shows a strong GC bias for all amino acid codons in humans. Sites not under significant constraint (D-F) do not show such a bias. For 4d sites under constraint, strong biases towards C or G bases largely reflect local base context, where sites that form CpGs are rare, as well as GGG codons that can lead to G-quadruplex structures in mRNA.

### Higher GC4 content in single exon genes and first exons

Mammalian transcripts typically contain introns. Under the UTH, native transcripts that are intronless should display high GC content at synonymous sites so that they are recognised as native transcripts despite lacking introns. We tested this by comparing the proportion of 4d sites that are GC across the mammal genomes in single- (n = 1,119) and multi-exon (n = 15,483) transcripts with >10 4d sites. Median GC4 in single-exon transcripts (0.67) is significantly greater than that in multi-exon transcripts (0.57; Wilcoxon rank sum test, W = 6,264,319, p < 2.2×10^-16^; Fig. 3A). As first and last exons are rarely spliced out^35^ it may be important that they maintain higher GC content as they are present in most transcripts and, as such, are most exposed to transcriptional silencing and degradation mechanisms, as well as containing essential splice sites. For exons with >10 4d sites, GC4 in the first exons of multi-exon transcripts are significantly higher than in other exons, (Fig. 3A; median GC4 in first exons = 0.72 (n = 6,070), other exons = 0.58 (n = 79,694); Wilcoxon rank sum test, W = 157,517,032, p < 2.2×10^-16^). Last exons also show significantly higher GC4 content, but the difference is smaller (median GC4 in last exons = 0.62 (n =8,053), other exons = 0.58 (n =78,830); Wilcoxon rank sum test, W = 287,613,287, p < 2.2×10^-16^). The length of exons may also relate to GC4 content, where higher GC content could be important in longer exons to signal them as native. However, we observe significant negative correlations between exon length and GC4 content for single exon genes (Pearson’s r = -0.18, d.f. = 1,117, p = 1.83×10^-9^) and in multi-exon transcripts (although the correlation here is extremely close to zero; Pearson’s r = -0.01, d.f. = 1,117, p = 7.2×10^-4^). We therefore do not find evidence for higher GC4 content in longer exons.

**Figure 3.**
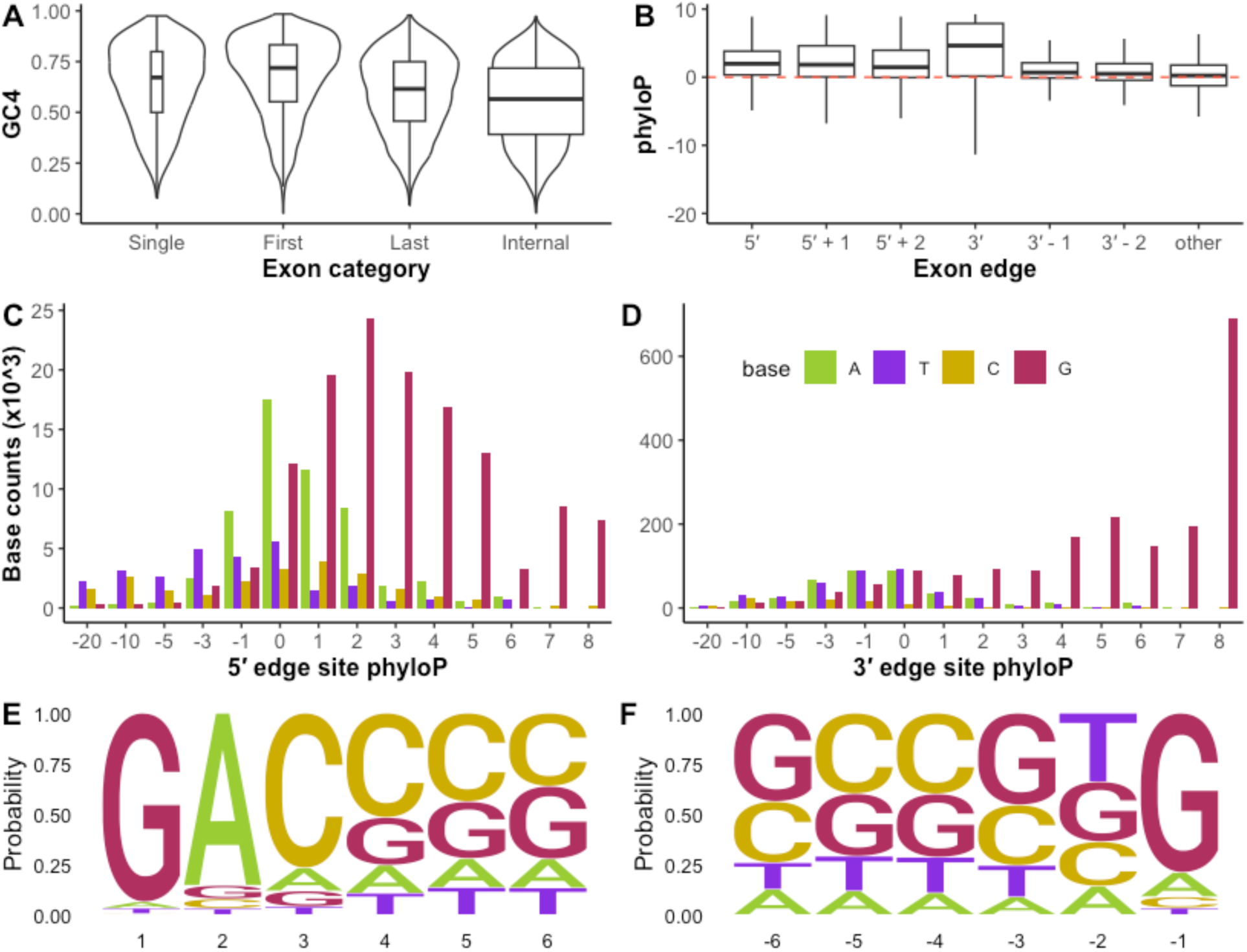
GC content of 4d sites in mammalian transcripts supports the unwanted transcript hypothesis. (A) GC4 content of single exon genes (n = 1,119) and the first (n = 6,070) and last (n = 8,053) exons of multi-exon genes are significantly greater than GC4 content of internal exons (Wilcoxon rank sum tests, p < 2.2×10^-16^ in all cases). (B) PhyloP is significantly higher at 4d sites within 3bp of exon-intron boundaries compared to 4d sites elsewhere in exons (ANOVA, F = 6,561, d.f. = 2, p < 2.2×10^-16^). Red dotted line indicates neutrality. Histograms of base counts across mammals at 4d sites at the first (5′ exon edge; C) and last (3′ exon edge; D) positions in exons in phyloP bins show strong bias for G at constrained 4d sites. Logo plots for base content of 4d sites at the first six positions (E) and last six positions (F) in exons, where the probability reflects the counts of each base across the mammal genomes at 4d sites located at those positions.

### High constraint at splice sites

The unwanted transcript hypothesis predicts high constraint at 4d sites involved in transcript splicing to avoid incorrect splicing that can lead to spurious transcripts. In line with this, we see significantly higher phyloP at 4d sites positioned at the end of exons (i.e., adjacent to splice sites) compared to 4d sites elsewhere in the coding sequence (Fig. 3B; n = 1,044 (5′ exon boundary), 12,049 (3′ exon boundary), 2,608,026 (elsewhere in the transcript); mean phyloP = 1.82 (5′ boundary), 3.70 (3′ boundary), -0.17 (elsewhere); ANOVA, F = 6,561, d.f. = 2, p < 2.2×10^-16^). A post-hoc Tukey’s test showed that constraint at exon 3′ boundaries is significantly greater than at 5′ boundaries (difference in mean phyloP = 1.88, p < 2.2×10^-16^). For 4d sites under constraint, there is a strong bias for 4d sites to be guanine at both 5′ and 3′ boundaries, with 79.1% and 93.6% G across the mammal genomes at these positions respectively (Fig. 3C,D). At the 5′ boundary, constrained 4d sites at the second position into exons (3′ splice site + 2 bp) show a strong bias for adenine (85.5%; Fig. 3D) and 10% of constrained 4d sites that are AT in the human genome are found at these positions (n = 10,359/103,961). Constrained 4d sites at the third position into exons at the 5′ end (3′ splice site + 2 bp) are mostly cytosine (76.8%; Fig. 3D). Overall, there is a strong bias for GC at constrained 4d sites close to exon boundaries, except at second positions inside exon-intron boundaries (Fig. 3E,F).

### Human variation reflects bias for GC at 4d sites

We used the TOPMed dataset of human genetic variation^33^, encompassing genome-wide single nucleotide polymorphism (SNP) allele frequencies from 138,922 individuals, to assess levels of variation at 4d sites in humans. Counts of singleton alleles (alleles present as a single copy in one heterozygous individual) in the TOPMed dataset reflect mutation bias expectations, with bias towards transition over transversion mutations and for CG to AT mutations. In particular, we observe a bias in CG to AT singleton transition mutations generally and at constrained sites (Fig. 4A,B). In the absence of selection a bias towards AT bases at 4d sites would be expected in humans. PhyloP scores of sites where singleton CG to AT alleles are observed are significantly higher than phyloP at sites with singleton AT to CG alleles (Fig. 4C; mean phyloP = 0.41 and - 1.45 at CG to AT and AT to CG sites respectively; t-test, t = -105.88, p<2.2×10^-16^). This reflects the fact that there are more CG 4d sites and that they are under greater constraint generally (Fig. 1D). However, at higher allele frequencies, CG to AT alleles are generally seen at sites with lower phyloP (mean phyloP = -2.42 and -3.51 at sites with low frequency (4×10^-6^ > MAF < 0.01) and common (0.01 > MAF < 0.5) CG to AT alleles respectively; ANOVA, F = 14,927, d.f. = 2, p<2.2×10^-16^). PhyloP at sites with low frequency and common CG to AT alleles is significantly less than phyloP at sites with low frequency and common AT to CG alleles (differences in mean phyloP between CG to AT and AT to CG alleles = -0.87 (low) and -0.61 (common); ANOVA, F = 23,448, d.f = 6, p<2.2×10^-16^). So, whilst CG to AT transition mutations are by far the most common seen in humans at 4d sites, they are under-represented at higher allele frequencies, particularly at sites under constraint (Fig. 4D,E).

**Figure 4.**
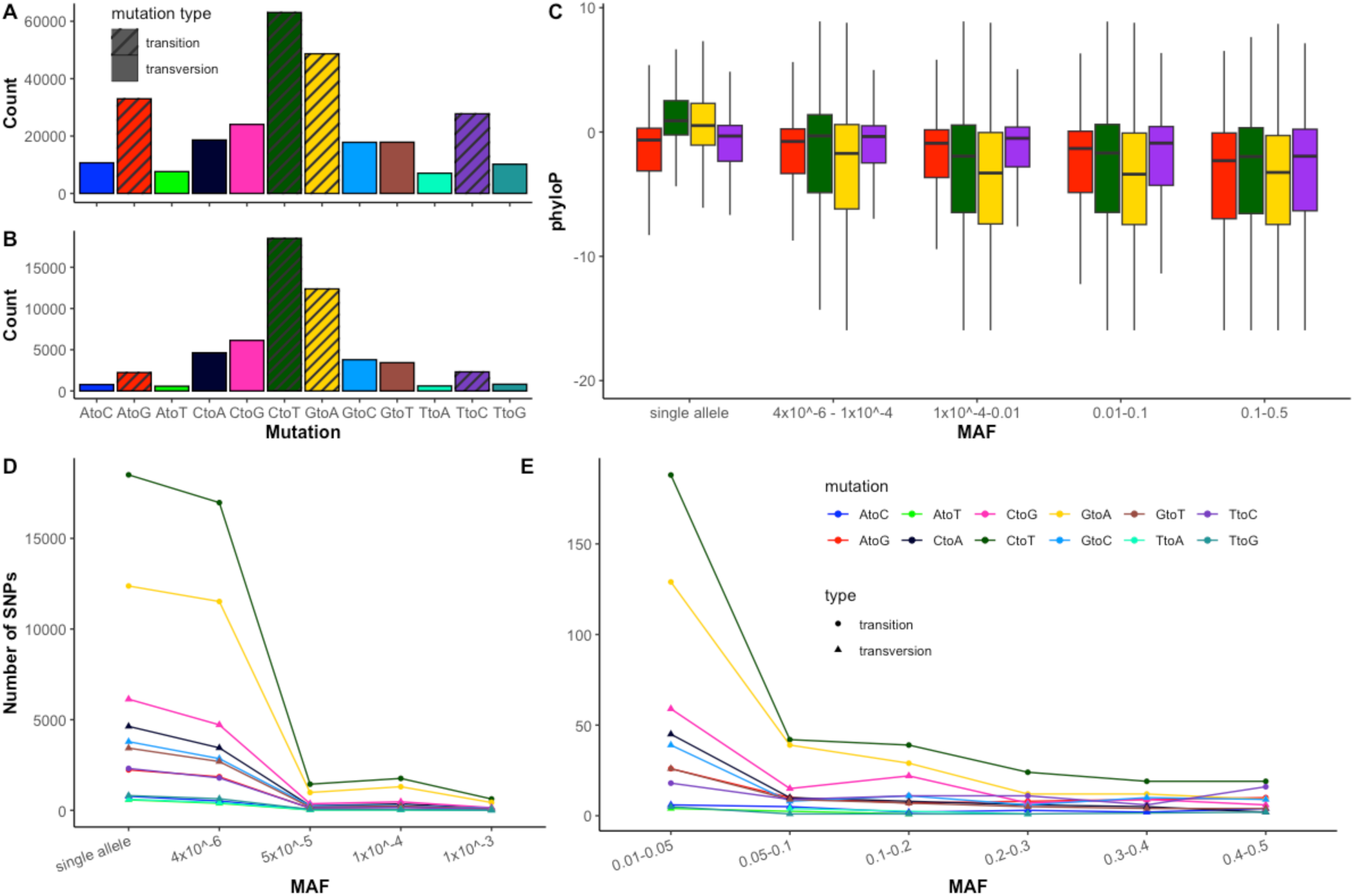
Human genetic variation at four-fold degenerate sites. Counts of singleton alleles present in only one individual across (A) all 4d sites and (B) constrained 4d sites show expected mutational bias, with more transition than transversion mutations and more CG to AT than AT to CG transitions. (C) CG sites with singleton transition SNPs are under greater constraint than those at AT sites, however this difference in constraint disappears and even reverses for SNPs with higher allele frequency, showing a strong selection pressure against CG to AT mutations at constrained sites. (D) Rare alleles are enriched for CG to AT mutations but, at higher allele frequencies (E), the bias for certain mutation types is greatly reduced.

### Associations between synonymous site variation, effective populations size, and genome transposable element content

The unwanted transcript hypothesis predicts that greater GC content of synonymous sites will be more important for species with low *Ne*, where inefficient selection can lead to a greater load of spurious transcripts^22^. We used mammalian *Ne* estimates generated as part of Zoonomia^36^ (n = 210 species) to test for a relationship between GC4 content and *Ne*. Mean *Ne* across the 210 species is 36,633 (s.d. = 39,034), with a ∼1,600 fold difference between the species with the highest *Ne* of 270,247 (prairie deer mouse, *Peromyscus maniculatus*, Order: Rodentia), and the lowest *Ne* of 162 (beluga whale, *Delphinapterus leucas,* Order: Cetartiodactyla). We do not see a significant correlation between *Ne* and GC4 (Pearson’s R = -0.13, p = 0.07). To assess whether *Ne* has an effect on GC content of highly constrained 4d sites, we reduced the data to a set of 4d sites that are represented in all 240 genomes (i.e., no missing data; n = 93,464 sites, mean phyloP = 3.84, s.d. = 1.32). We then estimated genetic distance among all species at these sites and generated a neighbour-joining tree (Fig. 5A). The tree recapitulates the topography of the expected placental mammal tree^37^, showing that divergence at constrained 4d sites is generally lineage-specific. GC content at these sites negatively correlates with historical *Ne* (Fig. 5B; Pearson’s R = -0.48, p = 5.3×10^-13^) as well as mean genetic distance to all other species (Fig. 5C; Pearson’s R = -0.87, p < 2.2×10^-16^). This demonstrates that constraint and GC content at these sites is lower in species with larger *Ne*, which is particularly apparent among rodents. The edible dormouse (*Glis glis*) is an outlier here, with exceptionally low *Ne* (*Ne* = 520) and the highest GC content at these constrained 4d sites among rodents (74%). The Indochinese shrew (*Crocidura indochinensis*, Order: Eulipotyphla) is the most divergent species, having the lowest GC content as well as the third highest *Ne* in the dataset (∼169,000). Contrastingly, Cetartiodactyla, particularly the cetaceans, are the most constrained at these sites, with among the highest GC content and lowest historical *Ne*.

**Figure 5.**
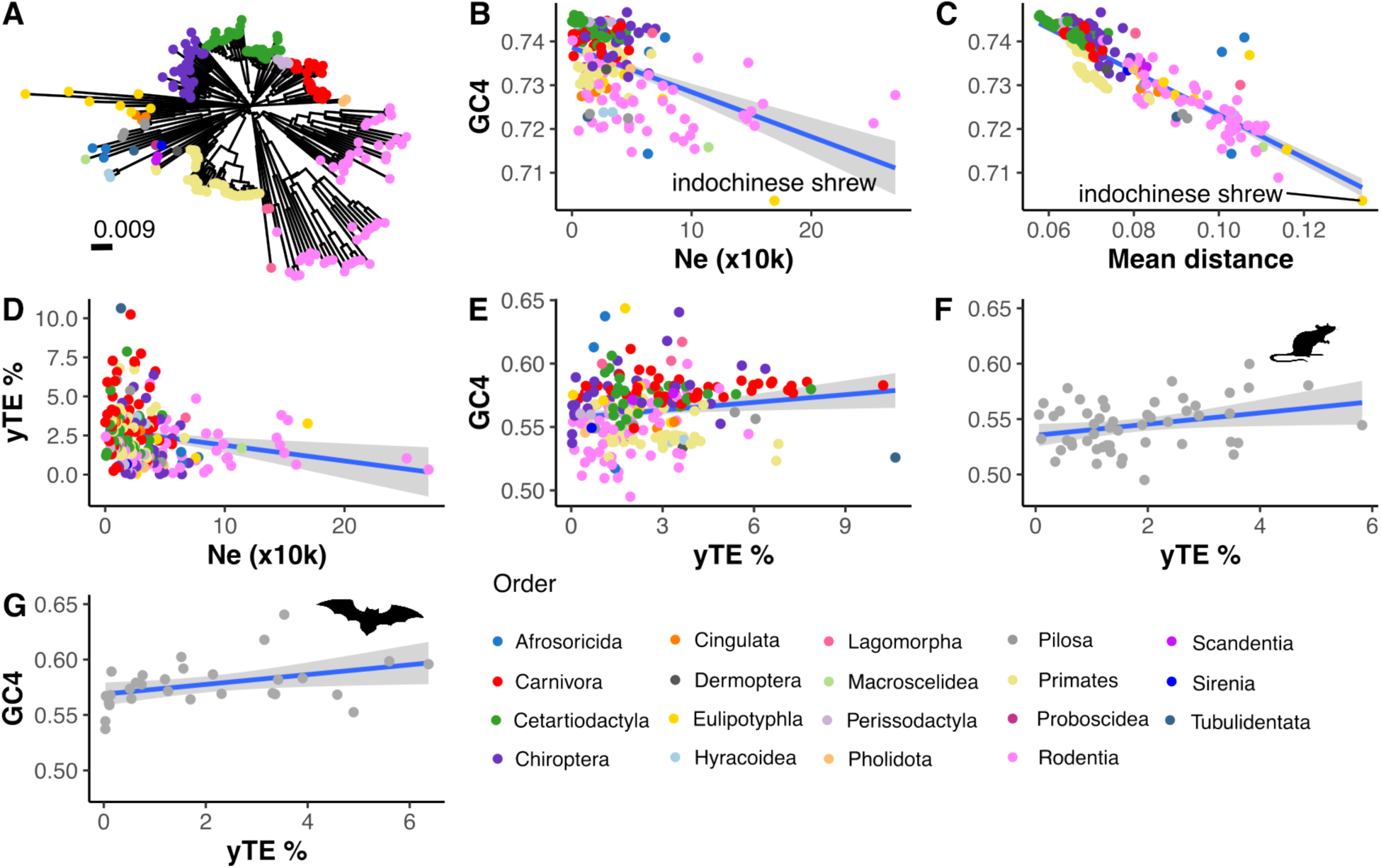
Effective population size and genome young transposable element content relates to GC content at highly constrained 4d sites. (A) Neighbour-joining tree of 240 mammals based on genetic distances (sequence dissimilarity) calculated from 93,464 shared 4d sites. Significant negative correlations are seen between GC4 content at highly constrained sites and (B) historical *Ne* (Pearson’s R = -0.48, p = 5.3×10^-13^) and (C) mean genetic distance of a species to all other species (Pearson’s R = -0.87, p < 2.2×10^-16^). Indicated in B and C is the Indochinese shrew (*Crocidura indochinensis*, Order: Eulipotyphla), which has the greatest mean distance to all other species, the lowest GC content at constrained 4d sites, and the third highest *Ne*. Genome young TE (yTE) content (a proxy for unwanted transcript burden) (D) negatively correlates with historical *Ne* (Pearson’s R = -0.20, p = 0.0034) and (E) positively correlates with GC content at 4d sites (Pearson’s R = -0.16, p = 0.016). The two orders with the strongest correlations between yTE content and GC4 content are (F) Rodentia (n = 53, Pearson’s R = 0.29, p = 0.035) and (G) Chiroptera (n = 30, Pearson’s R = 0.40, p = 0.031). Trend lines in B-G were calculated using a linear model with standard error intervals shown in grey.

Genome transposable element content may indicate (1) the efficiency of selection to purge unwanted/non-native sequences and (2) the burden of unwanted transcripts on a genome. A lower effective population size may therefore allow for higher genome TE content^24^ (although see^38^), as well as a need for higher synonymous site GC content, as the problem of unwanted transcripts may be greater in these species. We do not see a significant negative correlation between genome TE content and *Ne* (Pearson’s R = -0.10, p = 0.14). However, when looking only at young TEs (based on low divergence from consensus sequences^39^), which may be more indicative of the burden of TE-induced unwanted transcripts in a genome, the correlation is significant (Fig. 5D; Pearson’s R = -0.20, p = 0.0034). There is also a significant but weak positive correlation between genome young TE content and GC4 content across all 4d sites (Fig. 5E; Pearson’s R = 0.16, p = 0.016). We also looked for associations within orders, as levels of young TE content vary considerably among and within mammalian orders^39^. Both Rodentia (Fig. 5F; n = 53, Pearson’s R = 0.29, p = 0.035) and Chiroptera (Fig. 5G; n = 30, Pearson’s R = 0.40, p = 0.031) show significant positive correlations between young TE genome content and GC4 content.

### Distribution of constrained 4d sites among genes

Across the 17,394 transcripts within which we identified high confidence 4d sites shared among mammals, 16,860 (96.9%) of them contain at least one constrained 4d site and 12,257 transcripts (70.5%) have 10 or more constrained 4d sites. Excluding sites within 3 bp of exon boundaries, where constraint is generally high to ensure splicing accuracy, there are still 11,832 transcripts (68.0%) with 10 or more constrained 4d sites. This demonstrates how ubiquitous constraint at 4d sites is in mammal genomes. GC4 content can relate to the local GC content due to neutral processes such as GC-biased gene conversion (gBGC). We do observe a positive correlation between transcript GC4 content across mammals and local GC content (measured in 1Mb windows across the human genome; Fig. S1A; Pearson’s R = 0.52, p < 2.2×10^-16^). However, we observe a negative correlation between local GC content and transcript 4d site mean phyloP (Fig. S1B; Pearson’s r = -0.22, p < 2.2×10^-16^), showing that high 4d constraint is not associated with being in GC-rich genomic regions.

The top 1% most highly constrained 4d sites (phyloP ≥ 7.13, n = 26,282, GC4 = 99%) are distributed over 7,893 transcripts (45.4%), with 4,475 transcripts (25.7%) having at least two and 503 transcripts (2.9%) at least 10 highly constrained 4d sites. The gene containing the most top 1% highly constrained 4d sites is *TNRC6B* with 113 (13.1% of its 865 4d sites). Within this gene, 60% of its 4d sites are under significant constraint (n = 519, mean phyloP = 5.63, GC4 = 54.9%), with 72 sites that are fixed for the same base among all aligned genomes (GC4 = 61%). We assessed enrichment for constraint by comparing observed and expected rates of constrained 4d sites (Fig. 6A). There are 7,197 transcripts (43.4%) with observed/expected rates greater than 1, demonstrating that 4d site constraint is not distributed evenly among transcripts. The top 95th percentile of constraint enrichment (observed/expected rate > 1.98) contains 853 transcripts, and 166 transcripts are in the top 99th percentile (observed/expected rate > 2.83; Table S1). We also calculated enrichment of constrained 4d sites in CpGs and observe a subset of genes enriched for high 4d constraint in CpGs (Fig. 6B). The top 99th percentile (observed/expected ratio > 9.00) contains 115 genes (Table S1), 41 of which are also in the top 99th percentile of genes enriched for 4d constraint generally. Enrichment for constrained 4d sites is not explained by background GC content, with a lack of constraint observed at GC extremes (Fig. 6C,D). Constraint at CpG sites is also unrelated to being within CpG islands, with only five genes containing constrained 4d CpG sites being located within human CpG islands.

**Figure 6.**
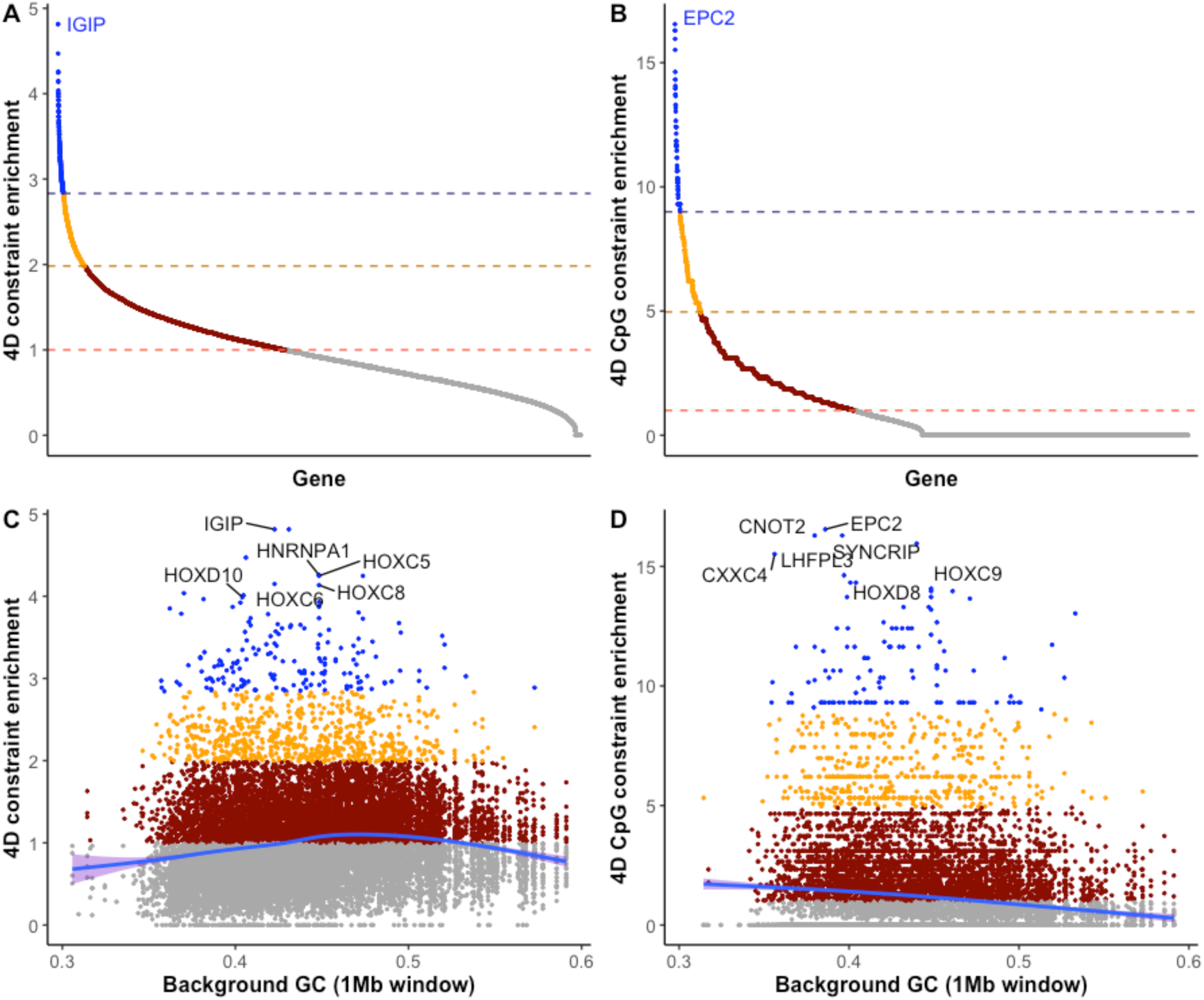
Genes enriched for constraint at four-fold degenerate sites are involved in development and transcriptional regulation. We estimated enrichment for constrained 4d sites (A) and for constrained 4d sites within CpGs (B) by comparing the observed to expected rate of constrained sites in each case (see methods). Genes with ratios >1 show evidence for enrichment, whereas those with ratios <1 (grey) have fewer than expected constrained 4d sites. The top 95th (orange and blue) and 99th (blue) percentiles of enrichment are indicated. Constraint enrichment is not explained by background GC content, with little enrichment observed in genes positioned in genomic regions with low or high GC. There is a slight positive correlation between background GC content and 4d site constraint enrichment (C; Pearson’s R = 0.10, p < 2.2×10^-16^) and a slight negative correlation between background GC content and 4d site CpG constraint enrichment (D; Pearson’s r = -0.15, p < 2.2×10^-16^). Smoothed trend lines in (C) and (D) were calculated using a general additive model with standard error intervals shown in purple.

Gene ontology analysis revealed strong enrichment for functions relating to development and transcriptional regulation in both sets (Tables S2,S3). The gene with the highest enrichment for 4d site constraint was *IGIP*, a single exon gene coding for immunoglobulin A inducing protein (17/17 4d sites under constraint). In the CpG enriched constraint set, *EPC2* (Enhancer of polycomb homolog 2) had the highest enrichment with 8 out of 9 4d sites in CpGs under constraint (enrichment score = 16.6). *EPC2* is predicted to be involved in transcriptional regulation and contribute to histone acetyltransferase activity*. CNOT2* (CCR4-NOT transcription complex subunit 2) also shows high enrichment for constrained CpG sites (enrichment score = 16.3). The CCR4-NOT complex interacts with *TNRC6B,* the gene with the most strongly constrained 4d sites, to regulate mRNA synthesis and degradation. Sixteen out of the 39 mammalian *HOX* genes (41%) are within this top 99th percentile of 4d site constraint enrichment, and 24/39 (62%) are within the top 95th percentile. *HOX* genes also show high prevalence of constrained 4d CpG sites, with 14 in the top 99th percentile of enriched genes. For example, *HOXC9* has 54 4d sites within CpGs, 40 of which are under constraint.

### Does constraint at 4d sites relate to overlapping regulatory features?

We performed a general linear model between 4d site phyloP and 15 variables (table 2) for all 4d sites with a phyloP ≥ 0 (n = 1,663,871) to test how much of the observed constraint can be explained by these variables. We removed all sites with negative phyloP scores as we were interested in testing for predictors of constraint (positive phyloP scores) rather than predictors of faster evolutionary rates (negative phyloP scores). The model explained 9.8% of the variation in phyloP and all but one variable (overlap with microRNA binding sites, p = 0.064) significantly explained some of the variance at p < 0.05. The variable that explained most of the variance, with a relative importance of 84.9%, was the GC proportion across mammals at each site. Distance to exon boundary (2.9% relative importance) and the background GC content (GC proportion of 1 Mb neighbouring sequence; 2.1% relative importance) were the next most important variables in the model. The remaining 11 variables had a combined relative importance of 10.1%, suggesting that while some synonymous sites constraint can be explained by being within regulatory features such as transcription factor binding sites, candidate *cis*-regulatory elements (cCREs), RNA binding protein (RBP) binding sites, or exonic splicing enhancers (ESEs), a large majority is explained by the base content of the sites, adding support to the unwanted transcript hypothesis.

**Table 2.** Outputs from a general linear model of synonymous site constraint in mammals. Model r^2^ = 0.098.

| Variable | Estimate | Std. Error | t value | Pr(>\|t\|) | Relative |
| --- | --- | --- | --- | --- | --- |

|  |  |  |  |  | importance |
| --- | --- | --- | --- | --- | --- |
| Intercept | -0.78 | 0.066 | -11.75 | $2.2 \times 10^{-16}$ | |
| GC proportion | 1.46 | 0.0039 | 372.15 | $2.2 \times 10^{-16}$ | 0.85 |
| Distance to exon edge | -0.00037 | $5.684 \times 10^{-6}$ | -65.61 | $2.2 \times 10^{-16}$ | 0.029 |
| Background GC proportion | -1.70 | 0.033 | -51.82 | $2.2 \times 10^{-16}$ | 0.021 |
| Within cCRE | 0.15 | 0.0039 | 40.02 | $2.2 \times 10^{-16}$ | 0.021 |
| CpG site | -0.51 | 0.0081 | -62.35 | $2.2 \times 10^{-16}$ | 0.015 |
| Within TFBS | 0.14 | 0.0045 | 30.34 | $2.2 \times 10^{-16}$ | 0.015 |
| Within uORF | 0.013 | 0.00030 | 43.92 | $2.2 \times 10^{-16}$ | 0.014 |
| Number of exons | -0.0052 | 0.00014 | -35.70 | $2.2 \times 10^{-16}$ | 0.013 |
| Number of aligned species | 0.011 | 0.00026 | 41.87 | $2.2 \times 10^{-16}$ | 0.0067 |
| Synonymous site $\mu$ | 4088.69 | 155.54 | 26.29 | $2.2 \times 10^{-16}$ | 0.0062 |
| Within ESE | 0.18 | 0.0047 | 38.74 | $2.2 \times 10^{-16}$ | 0.0048 |
| Within RBP binding site | 0.0034 | 0.00012 | 28.76 | $2.2 \times 10^{-16}$ | 0.0028 |
| Single exon | -0.16 | 0.0079 | -20.58 | $2.2 \times 10^{-16}$ | 0.0021 |
| Within lncRNA | 0.029 | 0.0050 | 5.80 | $6.78 \times 10^{-9}$ | 0.00041 |
| Within miRNA binding sites | -0.0053 | 0.0029 | -1.85 | 0.0639 | NA |

### Binding of certain transcription factors associates with 4d constraint

We tested whether there are certain TFs with binding sites that are enriched for 4d constraint, with the hypothesis that particular TFs may be involved in the regulation of genes that have high 4d constraint. We measured overlap between 4d sites and ENCODE 3 TF ChIP-seq clusters^40^ representing 340 TFs from 129 cell types. ChIP-seq signals are generally broader than the specific binding site and so overlap does not necessarily mean that a 4d site is in the specific binding sites of the TF itself^41^. Regardless, the observation of certain TFs being enriched for binding to or near highly constrained 4d sites can identify common regulatory features of the TF-bound genes. We found that the majority of TFs show no enrichment for binding to constrained 4d sites, with 60% of the TFs having an enrichment score below 0.01 and 95% an enrichment score below 0.59 (where 0 = no enrichment; Fig. S2A). However, four genes showed strong enrichment for constrained 4d sites in their TFBS ChIP signals, (99th percentile, enrichment score > 2.1), namely *EZH2* (enrichment score = 4.75)*, SUZ12* (enrichment score = 3.78)*, RNF2* (enrichment score = 2.36), and *POLR2A* (enrichment score = 2.24). The top three TFs (*EZH2, SUZ12, and RNF2*) encode proteins that are part of the Polycomb-group family involved in many cellular memory processes, including cell fate decisions during embryogenesis and maintaining the transcriptional repressive state of genes over successive cell generations^42–44^. There is a positive correlation between the number of TF binding sites overlapping constrained 4d sites for these three TFs and the transcript 4d constraint enrichment scores (Fig. S2B; Pearson’s R = 0.29, p < 2.2×10^-16^). This demonstrates that many of the genes enriched for 4d constraint, including several *HOX* genes, are bound by Polycomb-group repressors at or near constrained 4d sites. Genes in the top 99th percentile of 4d site - polycomb-group TF binding site overlap (n=168 genes; Table S4) are enriched for GO terms relating to transcriptional regulation and embryonic morphogenesis and development (Table S5). Genes absent of binding sites for the three polycomb group TFs but enriched for 4d site constraint (n = 109, Table S6) are enriched for GO terms relating to post-transcriptional regulation of gene expression (Table S7).

## Discussion

Whole genome sequencing and massive multi-species genome alignments are delivering unprecedented power and resolution for the characterisation of evolutionary constraint. Here, we have carried out a comprehensive characterisation of synonymous site constraint in mammalian genomes using the Zoonomia alignment of 240 placental mammal genomes in conjunction with single-base resolution constraint scores. We demonstrate a continuum of constraint at 4d sites and show that ∼21% are under significant constraint in mammals. Our estimate of synonymous site constraint is at the upper end of previous estimates in mammals, which range from 10-21.5%^7,45,46^. We argue that our comprehensive genome-wide analysis of constraint using single-base resolution constraint scores, provides the most complete picture to date of synonymous site constraint over the 100 million years of placental mammalian evolution.

We find several lines of evidence that constraint at synonymous sites may relate to reducing the issue of unwanted transcripts, in support of the UTH. We show a strong bias for GC4 content at constrained sites, which may help ‘flag’ transcripts as native^22^, and that single-exon genes and first exons of multi-exon genes have higher GC4 content across mammals, a pattern previously shown in humans^26^. We also show strong evidence for high constraint at 4d sites that occur at exon-intron boundaries, essential for ensuring the production of accurate transcripts^26,46–48^. Whilst constraint on 4d sites at exon boundaries is generally high, it explains <3% of the overall 4d constraint we observe. We find evidence for a negative relationship between GC4 content and *Ne*, as well as a positive relationship between GC4 content and genome young TE content as a proxy for unwanted transcript burden, both of which are predicted by the UTH.

The UTH does not claim to explain all constraint at synonymous sites and we show evidence for additional selective forces. In particular, we show strong enrichment of synonymous site constraint, including at CpG sites, in developmental genes and genes involved in transcriptional regulation. Genes enriched for 4d site constraint, such as *HOX* genes, are highly regulated genes and their accurate expression and timing of expression is integral to organism development and viability^49^. We suggest that the enriched 4d constraint likely relates to strong constraint on their regulation, such as methylation of CpG sites or targeting by transcription factors. Additionally, we show that some 4d constraint can be explained by having secondary functions, such as being within transcription factor binding sites, uORFs, or *cis*-regulatory elements.

Codon usage bias is a ubiquitous phenomenon in nature and all mammals show similar codon usage frequencies with a bias towards GC at codon third positions^50,51^. One hypothesis proposed to explain this is that the ancestral vertebrate genome was GC-rich, which would have created a prevalence of GC-ending codons early in vertebrate evolution^21^. Others have shown evidence for a substantial increase in GC3 content in certain mammalian lineages over the last 100 million years, likely related to differences in the strength of gBGC among species^52^, where AT:GC mismatches in recombinant sections are resolved with a preference towards GC^53^. We show here that GC4 content is variable among mammals, with a range from 49.5% in the hispid cotton rat (*Sigmodon hispidus*) to 64.4% in the European shrew (*Sorex araneus*). However, despite these differences which may be driven by gBGC, we demonstrate that ∼21% of 4d sites are under significant evolutionary constraint across the 240 genomes, with a strong GC bias. This GC bias is unlikely to be explained by gBGC, as a GC bias would also be expected to be seen in the surrounding sequence and not specific to the 4d sites. Whilst we do see a positive correlation between GC4 content of transcripts and local GC content, 4d site constraint in fact shows a slight negative correlation, with less 4d constraint observed in GC-rich regions. Additionally, recombination landscapes are expected to vary greatly across the genomes; recombination hotspots evolve rapidly^54^ with low conservation observed between humans and chimpanzees^55^, and between strains of mice local recombination rates correlate only very weakly^56^.

Maintenance of high GC content at the same sites is therefore most likely due to selection. At 4d sites not under significant constraint we do not observe a pronounced bias towards GC across mammals and, at sites with negative phyloP scores, we see a higher proportion of AT sites likely due to mutational bias. Our analysis of human genetic variation data revealed a strong bias in GC to AT mutations at 4d sites regardless of constraint which, in the absence of selection against them, should lead to a high AT content at 4d sites. However, GC sites under constraint lack higher frequency variants compared to AT sites, suggestive of purifying selection. This is in agreement with a recent analysis of selection against point mutations in the human genome, where no evidence was found for ultraselection (complete absence of point mutations) at 4d sites but 39% experience weak purifying selection^47^. Our lower estimate of 21% constraint is likely due to us characterising constraint across mammals, whereas the number of sites under constraint in any one species is likely to be considerably higher.

Hypermutability of methylated CpGs results in low CpG content of native transcripts generally. However, CpG content within the coding regions of certain genes is high. Intragenic methylation has been shown to protect gene bodies from aberrant entry of RNA polymerase II, which can lead to spurious expression^57,58^. We show that >16,000 4d sites that form CpGs are under high constraint in mammals. In particular, we see that certain developmental genes are enriched for constrained CpGs, including many *HOX* genes. These are small two-exon genes which overlap with regulatory CpG islands spanning from promoter regions to downstream of the coding regions. This thereby contributes to a substantial number of constrained CpGs in the coding sequence thought to be integral to their epigenetic regulation^20^. The constraint on developmental gene CpGs we demonstrate here suggests that these sites of epigenetic gene regulation via methylation have been maintained throughout mammalian evolution. Altered methylation of *HOX* genes has been shown to alter the development of mouse embryos^59^ and is associated with several diseases in humans^60^, including cancer^61^ and ageing^62,63^. Our characterization of constraint at CpG sites can therefore prove useful for dissecting the genetic and epigenetic causes of disease.

Two closely located (∼300 bp apart) single exon genes, *IGIP* and *PURA*, show amongst the highest enrichment of 4d constraint. *PURA*, which codes for pur-alpha, a short (322 amino acids) protein with highly conserved repeated nucleic acid binding domains^64^, has roles in controlling the cell cycle, DNA replication and transcription, as well as brain development^64^. These two genes are in a highly conserved region on chromosome 5, recently shown to be amongst the most invariable regions in humans^65^. Other developmental genes are also enriched for 4d site constraint, including many *HOX* genes, where highly orchestrated regulation of their expression during embryo development regulates regional character and patterning along the anterior-posterior axis^66^. Many of these genes are targets of the Polycomb-group TFs EZH2, SUZ12, and RNF2 and we see enrichment for constrained 4d sites in their ChIPseq binding site signals. As part of the Polycomb repressive complexes, these TFs bring about gene repression via epigenetic remodeling, are involved in cell fate decisions during embryogenesis, and their misregulation can lead to spurious transcript formation, leading to neoplastic transformation and cancer^67^. We see high enrichment of 4d constraint in several genes involved in transcriptional regulation, including *TNRC6B* whose protein localizes to mRNA-degrading cytoplasmic P bodies where it meditates miRNA-guided mRNA cleavage^68^. This gene works in conjunction with *CNOT2*, enriched for constraint at 4d CpG sites, which encodes a subunit of the CCR4-NOT complex, one of the major cellular mRNA deadenylases involved in bulk mRNA degradation, miRNA-mediated repression, and translational repression during translational initiation^69^. It is intriguing that a subset of genes involved in regulating gene expression processes, central to the UTH, show some of the highest constraint observed at 4d sites.

The UTH predicts that spurious transcription should be a greater issue for species with low *Ne* as selection is not efficient enough to remove weakly deleterious mutations that lead to spurious transcripts^22^. In support of this, we found that species with lower *Ne* generally have higher GC content at strongly constrained sites. In addition, *Ne* negatively correlates with, and GC4 content positively correlates with, the young transposable element content of a genome, which can be sources of spurious transcription. This is most strongly observed in bats and rodents, where variation in young TE content is highest. This evidence therefore suggests that mammals with smaller *Ne* experience greater selective constraint on their synonymous site content. Selection for error-mitigating properties in the error-prone genomes of species with small *Ne* has been previously suggested as an explanation for the observation of an association between *Ne* and intronic content and splice site usage across species^70^. Our findings support this hypothesis further, where higher GC4 content in species with lower *Ne* may act to limit the impact of spurious transcription and improve the detection of native transcripts amongst profuse unwanted transcripts.

*Ne* is generally associated with life history traits, making interpretation of these findings complicated. In mammals, species with low historic *Ne* are generally large and long lived, with low reproductive output. Short lived, high reproductive output species within the order Rodentia show the greatest variation in *Ne* in our dataset and many are among the species with the highest *Ne* and lowest GC4 content. The edible dormouse (*Glis glis*), the rodent with the lowest *Ne* and highest GC4 content in our dataset, has a unique life history among rodents, with a long life expectancy (average ∼ 9 years), a high yearly survival rate, and an infrequent breeding strategy due to an unpredictable food source^71^. The longest lived mammals in the dataset (cetaceans) show the highest constraint at their 4d sites among all the species. This link with longevity is intriguing; CpG methylation at or near *HOXL* subclass genes has been shown to be a strong predictor of longevity in mammals^63^ and changes within the mammalian chromatin landscape associate with aberrant transcription initiation inside genes during senescence and ageing^72^. Whether the high constraint on 4d sites in longer lived mammals suggested in our data reflects higher constraint on gene regulatory features, including intragenic CpG methylation of *HOXL* and other developmental genes, facilitates longer life by reducing the effects of spurious transcription will make for an intriguing follow up to this study.

## Methods

### Genome alignment and constraint scores

We used the 241-way multi-genome Cactus^73^ alignment of placental mammal genomes and human-referenced (Hg38) phyloP constraint scores generated as part of the Zoonomia project^30^. PhyloP scores were generated using the PHAST package^74,75^ with inputs of the Cactus alignment and ancestral repeats as a model of neutrally evolving sequences. PhyloP scores were generated at single-base resolution and ranged in value from -20 to 8.9. Positions under significant constraint were identified by converting phyloP values to *q* values using a FDR correction. Any positions with positive phyloP scores and a *q* ≤ 0.05 were considered to be under significant evolutionary constraint. This gave a significance threshold of phyloP ≥ 2.27 at 0.05 FDR. Further details can be found in the original paper^30^. PhyloP scores are available on the UCSC genome browser.

### Transcript selection and identification of mammalian 4d sites

Gene annotations for Hg38 were obtained from GENCODE release 39^31^. As most genes contain multiple transcripts we chose an approach to select one representative transcript per protein coding gene to simplify analyses. We selected representative transcripts using the Matched annotation from NCBI and EMBL-EBI set of representative transcripts.(MANE Select)^76^. If unavailable here, then we selected the canonical transcript used by gnomAD^77^ or BUSCO^78^. We used BEDtools intersect^79^ to extract phyloP scores for the CDS of each transcript from the single-base phyloP score files. Degeneracy of each site within CDS was assessed by looking up each codon in a DNA codon table using a custom perl script, revealing a total of 5,278,470 4d sites. Many of these are unlikely to be 4d sites in other mammal genomes due to, for example, differences in the gene content among genomes. We therefore extracted alignments for all codons from the Zoonomia alignment and identified codons where >85% of species were aligned and >95% of the aligned codons contained a 4d site (i.e. each 4d site is present in at least 194 species). This resulted in a set of 2,621,118 4d sites distributed over 17,394 transcripts that formed the basis of our analysis. Base content per 4d site per species were then extracted from the alignment and summarised using custom perl scripts. Whether a four-fold degenerate site forms a CpG site was determined from the human reference genome using a custom perl script which identified all 4d Cs followed by a G and all 4d Gs preceded by a C. Exon edge sites, exon counts per transcript, and single exon genes were identified from the GENCODE annotation. GC content in 1 Mb windows was measured across the human reference genome (GRCh38) using a custom perl script with a bed file as output. We then used the ‘intersect’ tool of BEDTools v.2.30.0^79^ between this file and a bed file of transcripts to obtain a local GC percentage for each transcript.

### Human population variation at 4d sites

We used the NHLBI Tans-omics for precision medicine (TOPMed) data freeze 8 (https://topmed.nhlbi.nih.gov/) containing whole genome sequencing of 138,922 individuals to assess human genetic variation at 4d sites. We used the ‘intersect’ tool of BEDTools v.2.30.0 to intersect the TOPMed VCF file and our 4d site bed file. At each 4d site with variants we ascertained whether or not the allele in the human reference genome (Hg38) matched the major allele in the human population using a custom perl script. This allowed us to identify variants that were specific to Hg38 and ensure that we were considering the most common allele when looking at base content of human 4d sites.

### Constraint enrichment

For each transcript we calculated the observed versus expected proportion of constrained 4d sites, constrained 4d sites in CpGs, and TFBS overlap with constrained 4d sites. In each case the expected proportion was calculated by 1/(total number of events/number of constrained events) (i.e. total 4d sites/total constrained 4d sites, total 4d CpG sites/total constrained 4d CpG sites, total TFBS-4d site overlaps/total TFBS-constrained 4d site overlaps). For each transcript, the observed rate was calculated by 1/(total events/constrained events). We then divided the observed by the expected proportion to get an enrichment score. We used the quantile function in R to identify highly enriched genes in each case, taking the top 95th and 99th quantiles. Gene ontology enrichment analysis was run on gene sets enriched for 4d constraint, 4d CpG constraint, and with constrained 4d sites enriched in TF binding sites using PANTHER^80^ with a Bonferroni correction for multiple testing.

### Effective population sizes, transposable elements, and divergence at 4d sites

Estimates of historical *Ne* for 210 of the Zoonomia species were obtained from Wilder *et al*., 2023, where they used PSMC^81^ to estimate *Ne* based on heterozygous positions in each genome. Data on genome transposable element content was obtained from Osmanski *et al.*, 2023, where they annotated the TE content of all Zoonomia genomes including identifying young insertions as TEs with sequences with K2P genetic distances < 4% compared to their consensus^39^.

To analyse divergence at highly constrained 4d sites we first identified a set of sites where all 240 species were aligned, giving 93,464 4d sites with no missing data . We then created concatenated pseudosequences in fasta format for each species from the alignment using a custom perl script. A single fasta format file containing these pseudosequences for all species was then read into R using the package Biostrings (DOI: 10.18129/B9.bioc.Biostrings). We then calculated a genetic distance matrix from the sequences using the R package DECIPHER (DOI: 10.18129/B9.bioc.DECIPHER), where each value in the matrix is the dissimilarity between two sequences. A neighbour-joining tree was then generated based on these distances using the Ape package in R (https://CRAN.R-project.org/package=ape). The tree was visualised using ggtree in R (DOI: 10.18129/B9.bioc.ggtree).

### Functional annotation datasets

Various datasets were used to look for relationships between constraint at 4d sites and functional annotations. All human cCREs were obtained from ENCODE SCREEN (https://screen.encodeproject.org/). Per-gene synonymous site mutation rates were obtained from supplementary dataset 11 in the gnomAD paper^77^. miRNA targets were obtained from TargetScanHuman release 8.0 (https://www.targetscan.org/vert_80/) and converted from hg19 to hg38 coordinates using the UCSC liftover tool (https://genome.ucsc.edu/cgi-bin/hgLiftOver). uORF coordinates were retrieved from http://github.gersteinlab.org/uORFs/^19^ and converted to hg38 using liftover. RNA binding protein binding sites were obtained from ENCODE eCLIP signals detailed in^82^ and available from https://www.encodeproject.org/. Transcription factor binding sites were obtained from ENCODE 3 TF ChIPseq datasets here: https://hgdownload.soe.ucsc.edu/goldenPath/hg38/encRegTfbsClustered/. Hg38 coordinates of constrained TFBs were provided by the authors of^30^. For identifying putative exonic splicing enhancers we downloaded the 238 hexamers identified as candidate ESEs in humans^83^ from http://hollywood.mit.edu/burgelab/rescue-ese/. We then used a custom perl script to check whether each 4d site was inside one of the hexamers to define whether a 4d site was inside or outside ESEs.

### Statistical analysis

All statistical analysis was carried out in R v.4.3.1 “Beagle Scouts”. For correlation analyses Pearson’s R was calculated, for comparisons between means two-tailed T-tests (two groups) or ANOVA (>2 groups) were used. Post-hoc Tukey’s HSD tests were used to identify significant differences following ANOVA. A general linear model (GLM) was used to look for associations between 4d site phyloP and multiple factors that may influence constraint.

## Acknowledgments

Computations and data handling were enabled by resources in projects NAISS 2023/5-90 and NAISS 2023/6-193 provided by the National Academic Infrastructure for Supercomputing in Sweden (NAISS), partially funded by the Swedish Research Council through grant agreement no. 2022-06725. KLT is a Distinguished professor funded by the Swedish Research Council.

## Author contributions

KLT and MJC conceived the project; MJC and MD carried out the bioinformatics; MJC performed the analyses; MD, JRSM, SVK and KLT contributed to interpretation; MJC wrote the first draft and all authors edited the manuscript.

## Competing interests

The authors have no competing interests.

## Materials and Correspondence

Correspondence and material requests should be addressed to M.J.C.

## Supplementary information

Supplementary tables file: Excel file containing Tables S1-S8.

Supplementary data file: Bed formatted file of the 2,621,118 mammalian 4d sites used in the study, with coordinates on the human reference genome (Hg38) as well as phyloP score and transcript ENSTXID for each site.

## Data availability

Scripts detailing all of the analyses are deposited on GitHub (https://github.com/MattChristmas/Mammalian_synonymous_site_constraint). The Cactus 241-way alignment and phyloP constraint scores are publicly available at https://cglgenomics.ucsc.edu/ data/cactus/ and at https://genome.ucsc.edu/cgi-bin/hgTrackUi?db=hg38&g=cons241way. Mammalian 4d sites that formed the basis of the analysis are provided as a supplementary data file in Hg38-referenced bed file format.

## Notes

### Competing Interest Statement

The authors have declared no competing interest.

