## Supplementary figures for "Interpreting mammalian evolutionary constraint at synonymous sites in light of the unwanted transcript hypothesis"

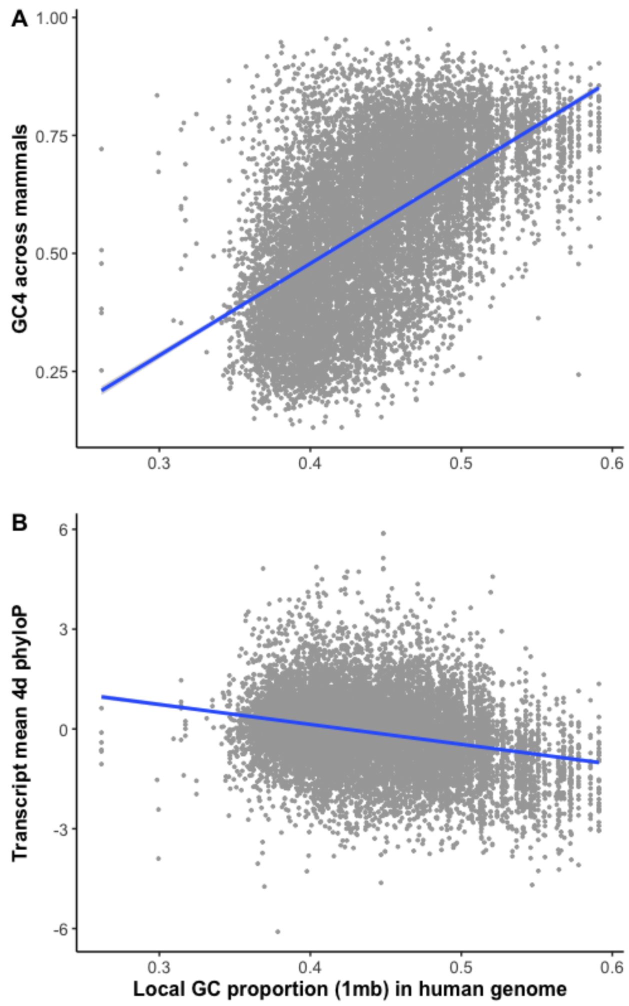


**Figure S1. Local GC content relates to GC4 content in mammals but cannot explain observed constraint** (A) Mammalian GC content at 4d sites per transcript significantly correlates with local GC content (1 Mb window) in the human genome (Pearson's $r = 0.52$, $p < 2.2 \times 10^{-16}$). (B) However, there is a negative correlation between local GC content and transcript 4d site mean phyloP (Pearson's $r = -0.22$, $p < 2.2 \times 10^{-16}$), showing that high 4d constraint is not explained by being in GC-rich regions of the genome generally.

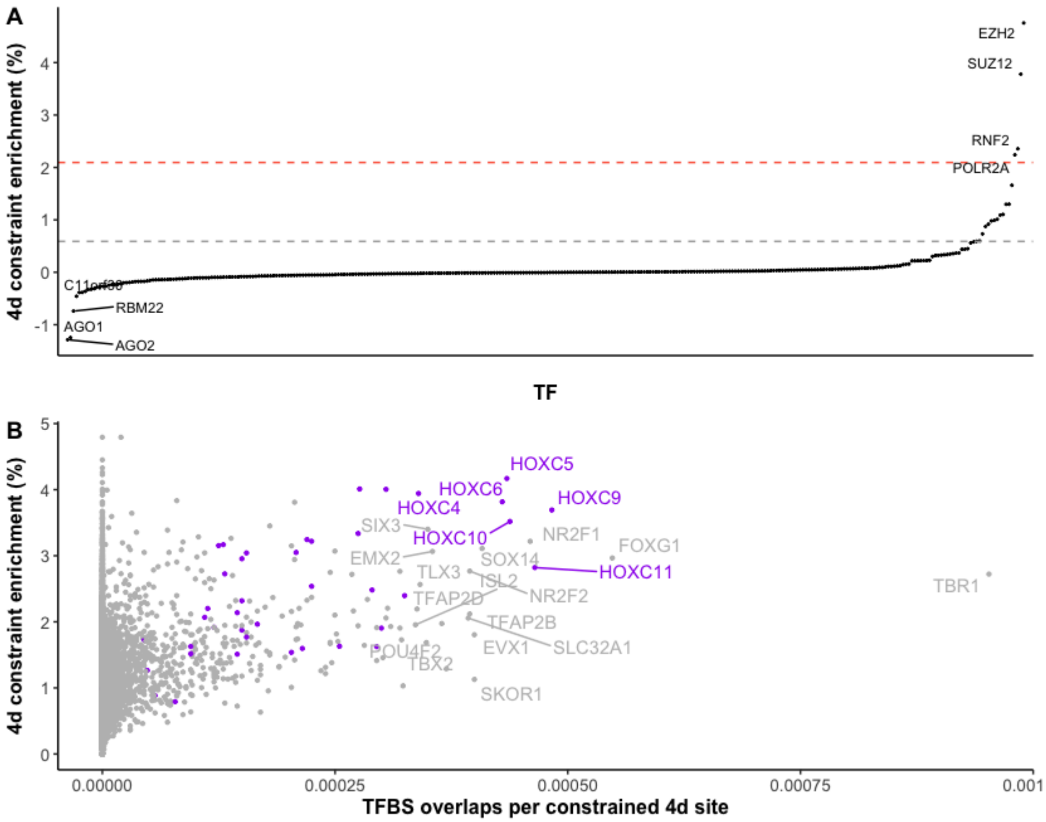


**Figure S2. Transcription factor binding sites and 4d constraint.** (A) Polycomb-group transcription factor binding sites are enriched for 4d site constraint. (B) Many developmental transcription factors enriched for 4d site constraint are bound by Polycomb-group TFs at or near their constrained 4d sites. *HOX* genes (purple) represent a group of developmental genes enriched for 4d site constraint, CpG constraint, and are bound by Polycomb-group TFs.
